# Free energy based high-resolution modeling of CTCF-mediated chromatin loops for human genome

**DOI:** 10.1101/105676

**Authors:** Wayne Dawson, Dariusz Plewczynski

**Affiliations:** Laboratory of Functional and Structural Genomics, Centre of New Technologies, University of Warsaw, Banacha 2c, Warsaw 02–089, Poland; Department of Biotechnology, Graduate School of Agricultural and Life Sciences, The University of Tokyo, 1-1-1 Yayoi, Bunkyo-ku, Tokyo 103-8657 Japan; Faculty of Pharmacy, Medical University of Warsaw, Banacha 1, 00-001 Warsaw, Poland; Centre for Innovative Research, Medical University of Bialystok, Białystok, Poland

## Abstract

In recent years, chromatin has been found to have considerable structural organization in the human genome with diverse parts of the chromatin interacting with each other to form what have been termed topologically associated domains (TADs). Chromatin interaction analysis by paired-end tag sequencing (ChIA-PET) is a recent protein-specific method that measures these chromatin interactions via specific interactions such as CTCF-cohesin binding proteins or RNA polymerase II interactions. Unlike high-throughput chromosome conformation capture (Hi-C), which measures unspecific binding (all against all), ChIA-PET measures specific protein-protein contact interactions; hence physical bonds that reflect binding free energies. In this work, a thermodynamic method for computing the stability and dynamics of chromatin loops is proposed. The CTCF-mediated interactions, as observed in ChIA-PET experiments for human B-lymphoblastoid cells, are evaluated in terms of a chain folding polymer model and the experimentally observed frequency of contacts within the chromatin regions. To estimate the optimal free energy and a Boltzmann distribution of suboptimal structures, the approach uses dynamic programming with methods to handle degeneracy and heuristics to compute parallel and antiparallel chain stems and pseudoknots. Moreover, multiple loops mediated by CTCF protein binding that connects together more than one chain into multimeric islands are simulated using the model. Based on the thermodynamic properties of those topological three-dimensional structures, we predict the correlation between the relative activity of chromatin loop and the Boltzmann probability, or the minimum free energy, depending also on its genomic length. The results show that segments of chromatin where the structures show a more stable minimum free energy (for a given genomic distance) tend to be inactive, whereas structures that have lower stability in the minimum free energy (with the same genomic distance) tend to be active.

## Introduction

In eukaryotic cells, detailed experimental identification of the structure of chromatin fiber inside of a cell has revealed considerable higher order organization and packaging in a hierarchical fashion (Barbieri et al., 2013a; Barbieri et al., 2013b; Davie, 1997; Davie et al., 2008; Dekker and Mirny, 2016; He et al., 2008; Lewis and Murrell, 2004; Lieberman-Aiden et al., 2009; Szalaj et al., 2016; Tang et al., 2015; Ulianov et al., 2016; Wang et al., 2015). Structural determination strategies have forged ahead with a number of high-throughput methods to obtain genome-wide maps of chromatin organization; e.g., chromatin interaction analysis by paired-end tag sequencing (ChIA-PET) and high-throughput chromosome conformation capture (Hi-C) (Dixon et al., 2012; Fullwood et al., 2009; Lieberman-Aiden et al., 2009).

The 3D topology of chromatin is thought to influence the regulation of gene expression and regions of active and repressed transcription in the cell, where diverse parts of the chromatin fiber can be found in proximity of each other in what are known as chromatin loops (Beagan et al., 2016; Buenrostro et al., 2015; Dekker and Mirny, 2016; Ji et al., 2016; Rao et al., 2014; Tang et al., 2015; Ulianov et al., 2016). In humans and other mammals, these loops are composed of insulator elements that bind in a complex composed of cohesin and the CTCF protein, an 11 zinc-fingers protein that binds CCCTC motifs (Ji et al., 2016; Rao et al., 2014; Tang et al., 2015; West et al., 2002). The CTCF forms dimers that can cause two distal regions of DNA to form into a loop (Ji et al., 2016). The RNA polymerase II (RNA pol II) complex is also strongly associated with the presence and formation of loops.

Recent work on chromatin structure has centered on including chromatin immunoprecipitation (ChIP) with ChIA-PET for specific protein factors (e.g., CTCF and RNA polymerase II) (Tang et al., 2015), offering structural information at a resolution of 1000 base pairs (1 kbp), given sufficient sequencing statistics. In general, a resolution of 5 kbp is now achievable as specific target proteins can be isolated - pushing whole genome structural resolution into the range of 200 nm and even smaller. A ChIA-PET experiment targets pairwise chain interactions (and multiple chain interactions) formed by a specific protein complex; in this work the binding of either CTCF or RNA pol II complexes. Loops often appear in the ChIA-PET data at specific, reproducible locations on the chromatin chain. Frequently observed pairwise interactions represent specific protein-protein binding interactions on the chromatin, and the strength of this interaction can be assessed based upon the frequency of the observed contact in the ChIA-PET data. Hence, ChIA-PET is largely impervious to unspecified binding such as self-ligation or random chain contact interactions. The frequency of observed contacts provides a measure of the binding proteins that form these loops and therefore are a function of the binding free energies of these contacts.

Unlike many protein and RNA structures where there is a very well defined 3D structure; chromatin structure tends to be far more plastic. Chromatin is dynamic; it can take on a variety of structures and thermodynamic states over time. Moreover, the interaction data typically is collected from around 100 million cells, each having a different threedimensional structure of the nucleus. Hence, the experimental ChIA-PET data reflects an ensemble of structures. Clustering of the structural ensemble yields a variety of meta-structures wherein the main structural feature is the pairwise interactions of different parts of the chromatin chain. These pairwise interactions between diverse parts of the chromatin chain via CTCF contacts (or RNA polymerase II) can be classified in terms of structural motifs. For uniquely definable interactions, a thermodynamic potential can be develop to estimate the likelihood of particular motifs within the ensemble.

For example, in CTCF structured motifs, the CTCF dimers combined with cohesin ring-like multi-domain structural proteins, bind in a parallel direction as a complex of proteins (Fig 1). The attachment to the chromatin chain can occur in an antiparallel or parallel direction at the loop anchor points - depending on the direction of the zinc fingers binding in the underlying DNA sequence motif. The most frequent and strongest interaction is the convergent loop (Fig 1A), where the chromatin chain is anchored in an antiparallel direction; this is shown in Fig 1A with the CTCF dimers and cohesin ring pointing in a parallel direction. Divergent loops (Rao et al., 2014) occur when the antiparallel chains have CTCF dimers anchored is in the opposite direction (Fig 1B). When the direction of the chromatin chain binds in a parallel interaction with the CTCF dimer, two types of structures of equal tendency and intermediate strength are suggested: tandem right (Fig 1C) and tandem left (Fig 1D) (Tang et al., 2015). In general, around 65% of the loops are convergent loops, roughly 33% are tandem loops and remaining divergent loops amount to only around 2% of the population (Tang et al., 2015). CTCF structures also appear to group into islands (as shown in Fig 1E), where the central region is the largely insoluble and inactive part (heterochromatin) whereas the regions jutting out are more accessible to transcription factors and therefore active (euchromatin).

**Figure 1.**
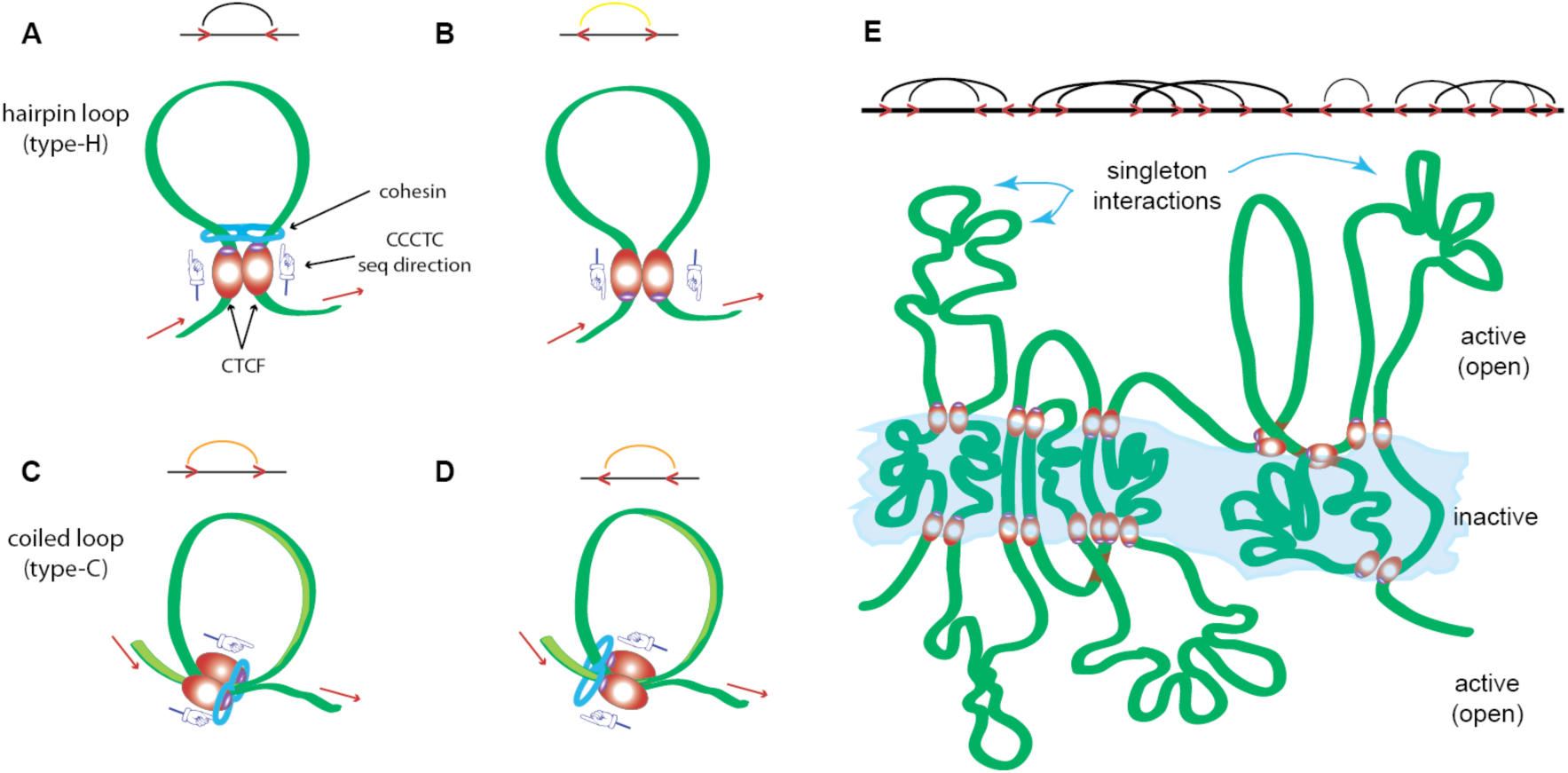
Examples of various types of loops formed by CTCF dimerization and its interactions with chromatin. (A) A convergent loop, (B) a divergent loop, (C) a tandem right loop, (D) a tandem left loop, and (E) combinations of A-D in the form of CTCF islands (a cartoon, but characteristic of regions like Chromosome 10). The 5’ to 3’ direction of the sense strand of the DNA sequence is indicated by the red arrows, cohesin is indicated by the blue feature enclosing structure, and the implied sequence direction for the CTCF motif is indicated by the directional pointers. The additional structural features are elaborated on in Supplementary Figs S1-S4.

The CTCF regions are the dominate structure motif in DNA regions containing insulators. However, within the long CTCF-mediated loops, often there are areas where RNA pol II is active. Such active regions tend to be spatially proximate to CTCF multimeric dense clusters, therefore the RNA pol II appears to be mediated by binding to CTCF-rich foci, active genes and enhancers; the RNA pol II loops appear to be smaller in general.

In addition to the CTCF interactions (large, stable loops) and RNA Pol II interactions (smaller and transient loops), there are singleton interactions: much weaker interactions that also involve CTCF-mediated binding or RNA Pol II binding (Tang et al., 2015). Singletons may sometimes involve divergent CTCF interactions (no cohesion, as in Fig 1B), sometimes RNA Pol II, still other times the effect of unknown transcription factors that interacts with CTCF or RNA pol II. When all these interactions are considered, many possible structural binding motifs are introduced: the CTCF dimers (Hirano, 2015; Tang et al., 2015) including various convergent and tandem loop motifs, the RNA Pol II regions spatially interacting (West et al., 2002), and the singleton interactions. A detailed categorization of loops and how they are evaluated in the program method are explained in the Supplement.

In recent years, there has been considerable interest in finding ways to model the distribution of chromatin structure physically using various approaches such as population-based analysis (Tjong et al., 2016) and polymer based models using molecular dynamics (MD) simulations (Barbieri et al., 2013a; Barbieri et al., 2013b; Chiariello et al., 2014; Meluzzi and Arya, 2013; Szalaj et al., 2016; Tang et al., 2015) or Monte Carlo (MC) simulation techniques (Wang et al., 2015). These models provide 3D structures of the chromatin. However, MD or MC simulation techniques do not guarantee an exhaustive search of such a large landscape for chromatin structural motifs, nor is it easy to say that a structure found in this way is actually the most stable structure (Tjong et al., 2016). More recently, a maximum entropy method has been introduced (Di Pierro et al., 2016; Zhang and Wolynes, 2017) that explores the phase space of the polymer chain under the constraints of pair interactions and uses these contacts to fit a proposed 3D potential using maximum entropy.

Since a principal structural feature of chromatin (over the length scale of some 3 Mbp) is its 2D meta-structure, it is worth employing a dynamic programming algorithm (DPA) strategy to optimize a given thermodynamic representation of the pair contacts. Moreover, chromatin tends to have a hierarchical loop structure (Bascom and Schlick, 2017) that suggests typical thermodynamic folding observed in other biopolymer. Therefore, we have adapted the DPA with some additional heuristic modifications to search for meta-structure motifs such as loops and stem-like patterns that can take on a relatively stable form in a very specific conformation. Exploring the 2D landscape of a set of meta-structures can be done quickly and efficiently using a DPA heuristic based on free energy. The experimentally observed pair interaction frequency (PIF) can be used to estimate the binding affinity or statistical weight for a given pair. A pair binding free energy based polymer model can be constructed to model motifs from a statistical potential using the PIFs (from ChIA-PET data) and entropy loss due to folding model that measures the entropy loss due to the formation of contacts, or formation of cross links ((Dawson et al., 2001a, b; Dawson et al., 2014; Dawson and Kawai, 2015) and references therein). This would inform us of the optimal structures of these loops, or collections of loops from chromatin capture domains (CCDs) and whether such a distribution is largely flexible and shifting between multiple structures (a likely case for active regions; e.g., related to the presence of RNA Pol II), or a single structure that is largely frozen (a likely case for an inactive region of the genome, enriched with insulators such as CTCF proteins).

Moreover, an important attribute of this free energy approach is that active and inactive or open and closed states of the CCDs are found to be consistent with a minimum free energy interpretation wherein a larger number of contacts and their statistical weights are correlated with inactive or closed regions when scaled with genomic distance. The stability of particular conformations and motifs in an ensemble of structures should correlate with the PIF and this, in turn, should also correlate with the free energy. A more negative free energy (scaled with sequence length) should show a strong correlation with inactive regions of the genome due to freezing out (slowing down) of the dynamics, and vice versa for the active regions. The free energy should thus reveal regions composed of euchromatin (more unpacked regions) and heterochromatin (regions that are rather densely packed).

Therefore, in this work, based upon the observed PIF of the chromatin fiber contacts obtained from ChIA-PET data, we have developed a free energy based model with suboptimal structures that extracts the major part of the Boltzmann distribution of the ensemble of chromatin structural motifs distinguished in terms of singletons, CTCF, RNA Pol II binding sites, and multimeric CTCF islands. The approach handles complex motifs such as pseudoknots (involving both parallel and antiparallel chain interactions - see Supplement) using heuristic adaptations to the DPA. This model works largely from the opposite direction of (Di Pierro et al., 2016). Here, we combine experimentally determined data for contacts with a quantitative measure of entropy loss in the chromatin chain due to the formation of contacts or cross links. The entropy loss in this model is a *bottom up* integration procedure, where the maximum entropy is the unfolded chromatin chain and the free energy of a given folded chromatin structure represents the competition between the entropy loss and the binding energy of the contacts (or cross links). We then, for purpose of demonstration, use this contact information to explore 3D conformation under restraints.

We stress that this is not intended to be a thorough study of everything has ever been done in Hi-C. We have used ChIA-PET data from a particular cell line, which is published elsewhere (Szalaj et al., 2016; Tang et al., 2015) and used this data on a whole genome to test this model as a proof of principle.

## Methods

The details of the methods are explained in the Supplement. Here we briefly explain the concepts.

### Thermodynamics and meta structure analysis algorithm

The model assumes a highly coarse-grained resolution ranging. Currently, we have tested resolutions ranging between 1 kbp and 20 kbp, where in this work, we focus on 5 kbp. At this scale, the standard measure of stiffness of a polymer, known as the persistence length or the Kuhn length (Hagerman, 1988), can be largely neglected because the persistence length is certainly less than 5 kbp; though it may possibly exceed 1 kbp in some cases. These meta-structures are treated as uniform beads on a string, though it should always be remembered that this is *not* how the real situation is at atomic resolution scales (Todolli et al., 2017). This is just a convenient approximation. Details are provided in the Supplement.

The computational approach is a thermodynamic optimization procedure, which employs adapted dynamic programing algorithm (DPA) procedures, to merge the entropy contributions of a Gaussian polymer chain (GPC) model (which expresses the polymer folding of chromatin) and the observed pair end tag (PET) counts from ChIA-PET experiments as statistical weights (which are converted to statistical potentials that compete against the folding entropy). A thermodynamic model permits a way to discern the principal structures of the chromatin found in the ensemble. The PET counts in ChIA-PET data represent inter-ligation of specific protein-protein binding of the CTCF complex. Hence, a result of this optimization procedure is the generation of meta-structures (ranked with respect to free energy) that express the pair-wise interactions of the chromatin chain as well as multi-chain interactions. The method has been demonstrated to work for other problems such as RNA structure prediction with pseudoknots (Dawson et al., 2007; Dawson et al., 2014) and in proof of principle in protein structure prediction (Dawson et al., 2005). This is the first time to test this GPC-based entropy model based on chromatin structures.

The program analyzes for the following structural motifs: hairpins, internal loops, multiloops, stems-like interactions, pseudoknot-like interactions and CTCF-islands. Heuristics in the DPA were introduced to handle pseudoknot-like interactions on the chain and islands of CTCF bound proteins that appear to involve multiple chain interactions. With these heuristics, the time complexity of the computational method is approximately *O(N* ^3^), where *N* is the number of beads in the chromatin sequence. After evaluating the free energy, a trace back procedure selects the principal structural motifs at the top of each pushdown stack and traces out the optimal solution using the connection information stored at the given motif. Suboptimal structures are built in a similar way by employing other combinations of solutions in the pushdown stacks. The details of this part of the algorithm can be found in the Supplement.

The package consists of various object oriented programs implemented using the python 2.7 platform. The 2D graphics of contacts are displayed using VARNA (Darty et al., 2009) and the 3D plots are generated using Gnuplot (Williams and Kelley, 2014).

### 3D structure construction

Meta-structures are much easier to understand visually than full 3D structures; nevertheless, the pairing contacts from these meta-structures can be employed as restraints in various MD simulation programs to generate a 3D ensemble of structures. As proof of principle, for the example results presented in this work, we show a representation of the 3D ensemble that was generated by replica exchange Monte Carlo simulation methods from the coarse grained package SimRNA (Boniecki et al., 2016), in which the histogram weight parameters were set to zero and a sequence of poly(A) was constructed and folded under the corresponding restraints generated for a given specified structure. The resulting trajectories can then be rescaled to the appropriate length scale of chromatin. The current fits are restricted to a range of a few Mbps; significantly smaller than the total length of the chromatin chain, about 3 Gbps. Hence, the closed boundary conditions of the cell nucleus can be neglected. On this range, the meta-structure represents the dominant structural features of the chromatin. Structures in this work are viewed using VMD (Humphrey et al., 1996).

### Acquisition of PET counts

The ChIA-PET data is obtained for CTCF contacts and RNA pol II information. It has an interaction range for self-ligated of about 8 kb (which are removed) and all other structures should be considered inter-ligated interactions (Szalaj et al., 2016). In this work, the data was binned into segments of length 5 kbps and divided into chromatin contact domain (CCDs) - loops or anchors of various sizes that were found in the experimental data. Within these CCDs, singleton, tandem loops, convergent loops and divergent loops where found - depending on their PET count. In the analysis, all PET counts greater than 4 are treated as anchors, where 4 and less are singletons. From these CCDs, heat-maps were generated that express the relative positions of contact (in 5 kbp segments) and the number of PET counts. In this work, the corresponding contacts were from previously published data using other methods. These heat-maps express an ensemble of particular chromatin structures where many of the pair contacts overlap.

## Results and Discussion

There are two objectives in this work. The first is to use polymer physics to obtain information on the dominant stable structural motifs of a given ensemble of observed contacts from a particular cell line (in this case, GM12878) and to discern the dynamics of the chromatin. The second is to use this information to identify regions of active euchromatin and inactive heterochromatin. The free energy helps to prioritize the types of motifs that dominate a particular region of the chromatin. The results presented here are intended as a proof of principle presentation of the model, not a model fully arrived.

The contact map data obtained from experiments provides us with an ensemble of the observed contacts (pairs) within the large population of cells. The thermodynamics of polymers provides us with an understanding of the influence of any given collection of contacts on the observed structure (or structures) in the ensemble in the form of a statistical weight; i.e., the Boltzmann distribution. The resulting structures are 2D because what is observed is not one specific 3D structure, but individual contacts between diverse parts of the chromatin fiber. Using thermodynamics distinguishes the relative contribution from different contacts and the likelihood of particular structures. This in turn permits some picture of the likely dynamics of chromatin from the experimental data using the thermodynamic probability of different structures.

In this model, we assume that the observed frequency of contacts is identical for every cell. Most likely, when millions of cells are measured, each particular cell will have common housekeeping genes where the expression is identical, but particular states of the cell that reflect that particular conditions of that given cell. The configuration of these particular states is largely unknown. However, since the CTCF contacts are the source of the major interactions, in any particular loop study, if there were truly an issue with different cell states in the population, it would be possible to turn on and off these interactions and fit the observed ensemble of cells. This will be addressed in future work.

When one particular structure dominates all the other structures by a substantial margin, this means that most of the time, the structure remains fixed with the given set of contacts and all the other configurations are occasional or incidental. This is a property that one would expect of heterochromatin, the densely packed regions of chromatin where very little expression occurs. This may be regions wherein the chromatin is not used for a particular part of the life cycle of the cell: such regions might include developmental genes in an adult, or regions of the chromatin that are only expressed when the cell is under stress. We would expect that a structure with a large negative free energy is indicative of very little dynamics; i.e., tightly packed. When only a few very similar structures dominate the distribution, we would assume from this ansatz that the chromatin is shifting between various states but is only somewhat dynamic. When there are many diverse structures that have rather similar probabilities, then we can assume that there is very little differentiation between such the regions of chromatin. This latter condition would be what is characteristic of euchromatin, regions where there is generally a lot of gene expression and where the structures are open and ready for transcription.

Therefore, the thermodynamic probability of a given structure relative to others should be a helpful measure of to what extent that particular region of chromatin is heterochromatin or euchromatin.

Fig 2 shows a relatively simple example of an observed heatmap for a small CCD from the CTCF data. Fig 2A is the experimentally observed heatmap (contact map), Fig 2B shows the dominant contacts found with the polymer model developed here, and Figs 2C show a 2D representation of the minimum free energy structure with the corresponding ensemble of 3D structures in Fig 2D. Many features of the original heat map are observed in the thermodynamic distribution, indicating a reasonable correspondence between the observed frequency of contacts and the actual distribution of structures that results.

**Figure 2.**
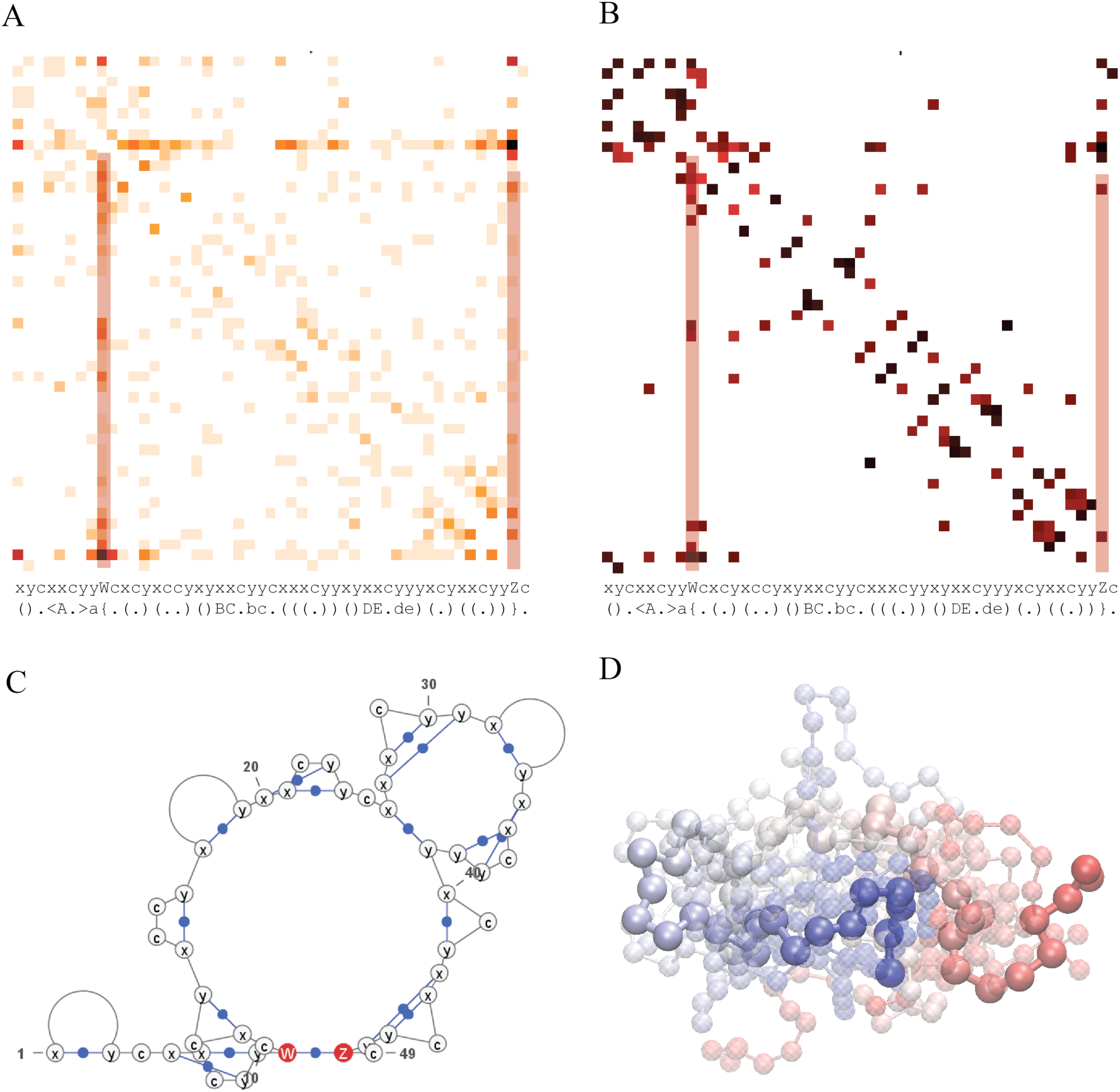
An example of contacts within a loop obtained from PET clusters and the weighted thermodynamic distribution of those structures based on the assumed polymer behavior of the chromatin. A) The original heat map of the data obtained from ChIA-PET data. B) The analysis using of predominant structures based on the Boltzmann distribution. C) The dominant structure found in the distribution displayed using VARNA. (D) The ensemble of 3D structures generated from the structure in panel C, displayed using VMD.

Fig 3 is a more complex structure showing both parallel strands and CTCF islands. Fig 3A shows the observed contact map along with the 1D structural notation, Fig 3B shows the dominant contacts found with the polymer model developed here with the sequence also shown for comparison, and Fig 3C and D shows a 2D representation of the minimum free energy structure and a corresponding ensemble of 3D structures, respectively.

**Figure 3.**
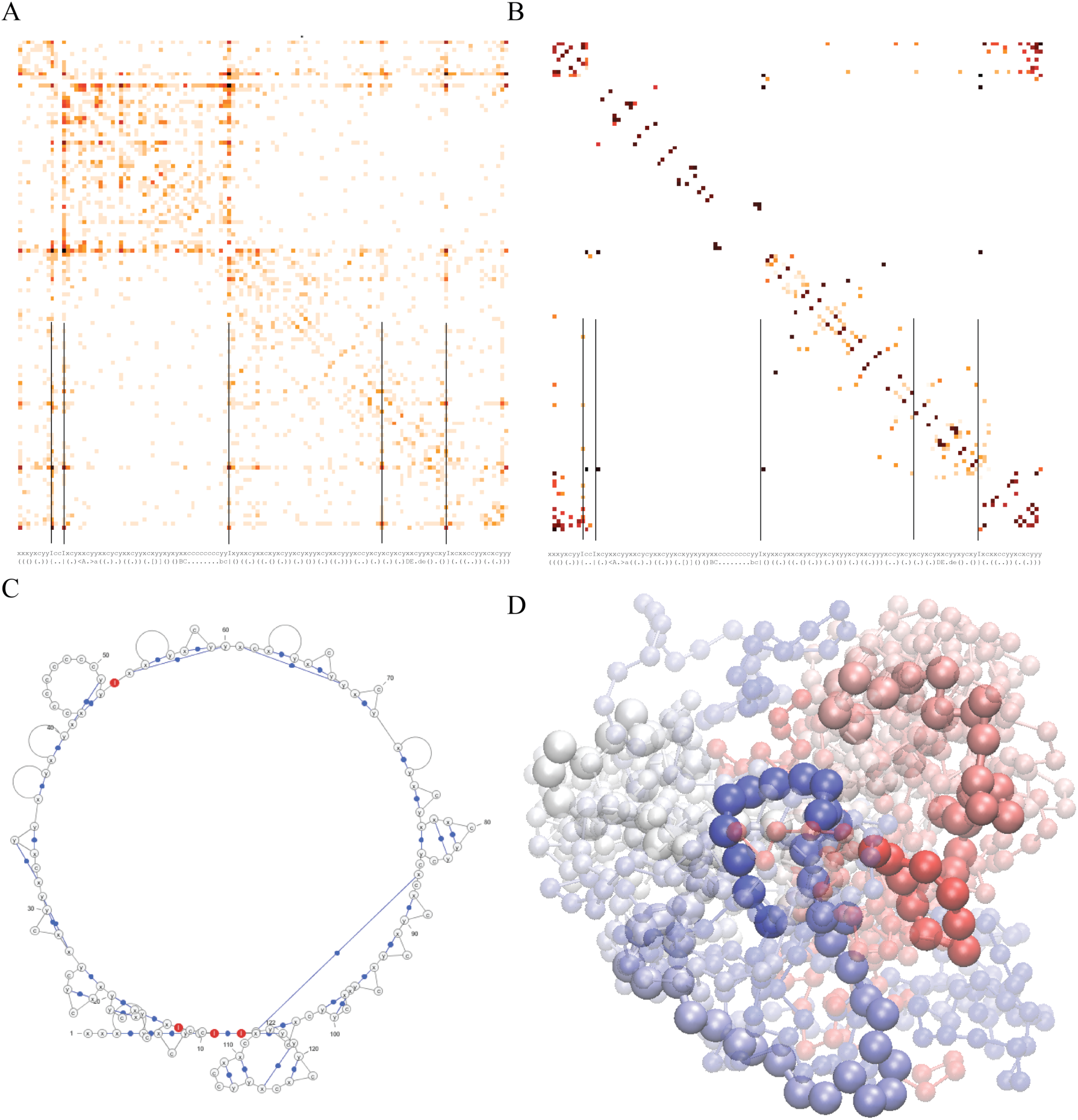
Result of A more complex structure in which the structure form CTCF-islands. This also shows an example of contacts within a loop obtained from PET clusters and the weighted thermodynamic distribution of those structures based on the assumed polymer behavior of the chromatin. A) The original heat map of the data obtained from ChIA-PET data. B) The analysis using of predominant structures based on the Boltzmann distribution. C) The dominant structure found in the distribution displayed using VARNA. (D) The ensemble of 3D structures generated from the structure in panel C, displayed using VMD.

This general correspondence permits us a means to examine the condition of the chromatin in the nucleus of the cell. Therefore, we calculated the minimum free energy structures and major suboptimal structures from all the available CCDs generated in previously published study (Szalaj et al., 2016; Tang et al., 2015) and plotted it in terms of the free energy, genomic distance and the experimentally established degree of activity and openness. The data for whether the structure is active or inactive and whether it is open or closed is obtained independently based on epigenetic markers. Hence the active/inactive axis or open/closed axis is independent of these free energy calculations.

In this perspective, it is assumed that regions that are inactive have very stable structures - the structure with the minimum free energy (mFE) is significantly more negative (for a given genomic distance) and the distribution of alternative structures of similar free energy are few. This means there is little chance that such chromatin will be found unpacked. Likewise, if the region is active, then the mFE structure is one of many of similar probability. This would tend to be a consequence of the chromatin being dynamic, with no particular structure heavily dominating the ensemble.

Based upon the published CCD data for loops over the whole genome (Szalaj et al., 2016; Tang et al., 2015), we evaluated minimum free energy (mFE). From the mFE, we also analyzed whole genome for the activity of the chromatin. Figs 4A and C show that when the active regions are plotted as a function of genomic distance and mFE, the active regions generally have a less negative FE compared to the inactive regions; indeed the behavior is curiously stable. Fig 4B and D show the measure of openness and correlates with the activity in Figs 4A and C. Regions that are largely inactive tend to have very stable structures where the mFE structure in the Boltzmann distribution is significantly more negative. The Boltzmann probability of the optimal structure appears to be mainly a function of the genomic length and only mildly influenced by the activity (Fig 5). The main tendency is the expected decrease in probability with genomic distance. This is because a longer segment of chromatin has many more possible states than a shorter segment.

**Figure 4.**
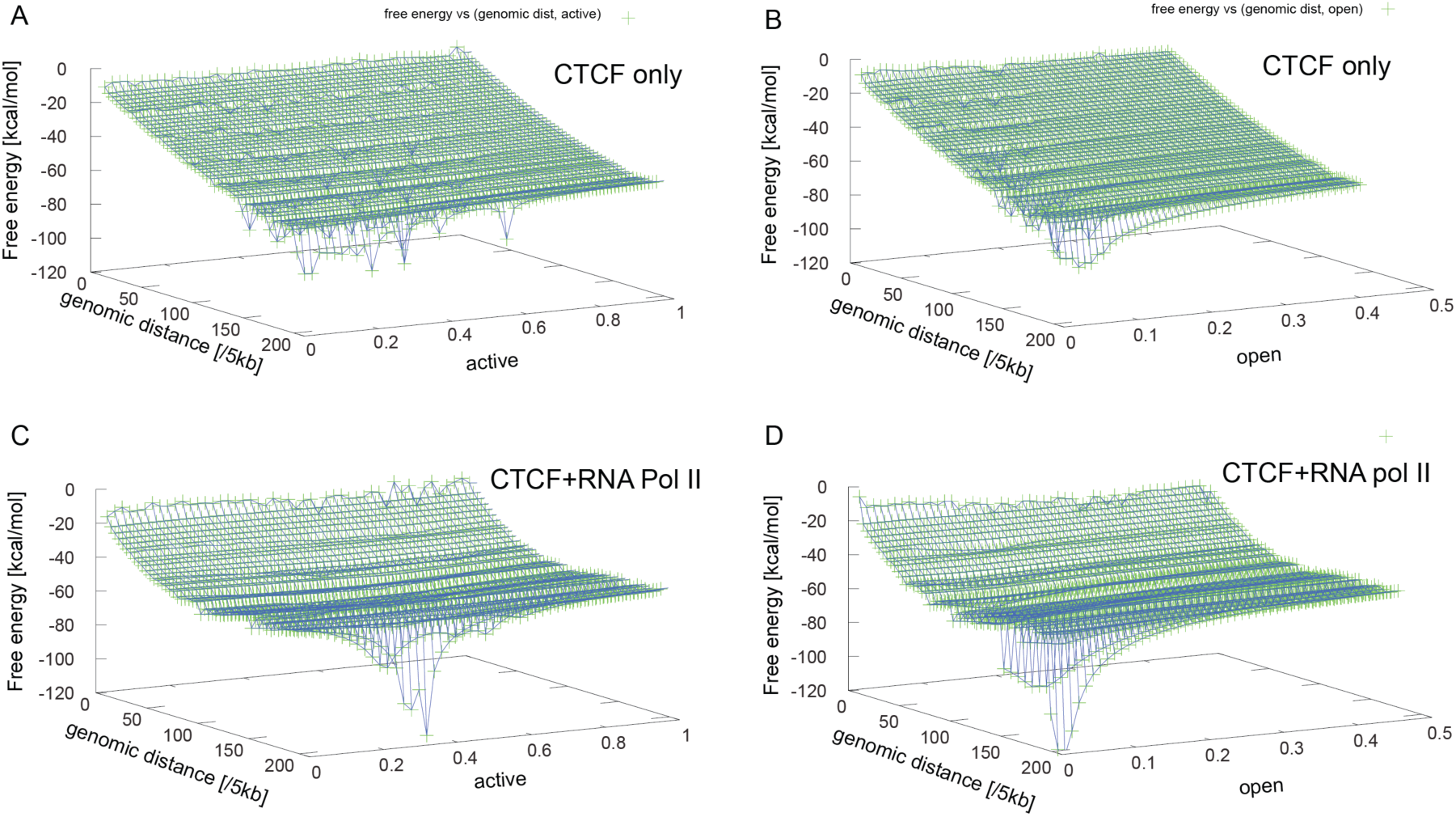
Comparison of the minimum free energy plotted with respect to both the genomic distance and the degree of activity or openness for CTCF and RNA Pol II. A) and B) Respective activity and openness for CTCF binding targets. C) and D) Respective activity and openness for CTCF with RNA pol II binding targets. (B) loops. For the same genomic distance, structures that have a more negative free energy show a tendency to be inactive and closed, and structures that are less negative tend to be more active and open.

**Figure 5.**
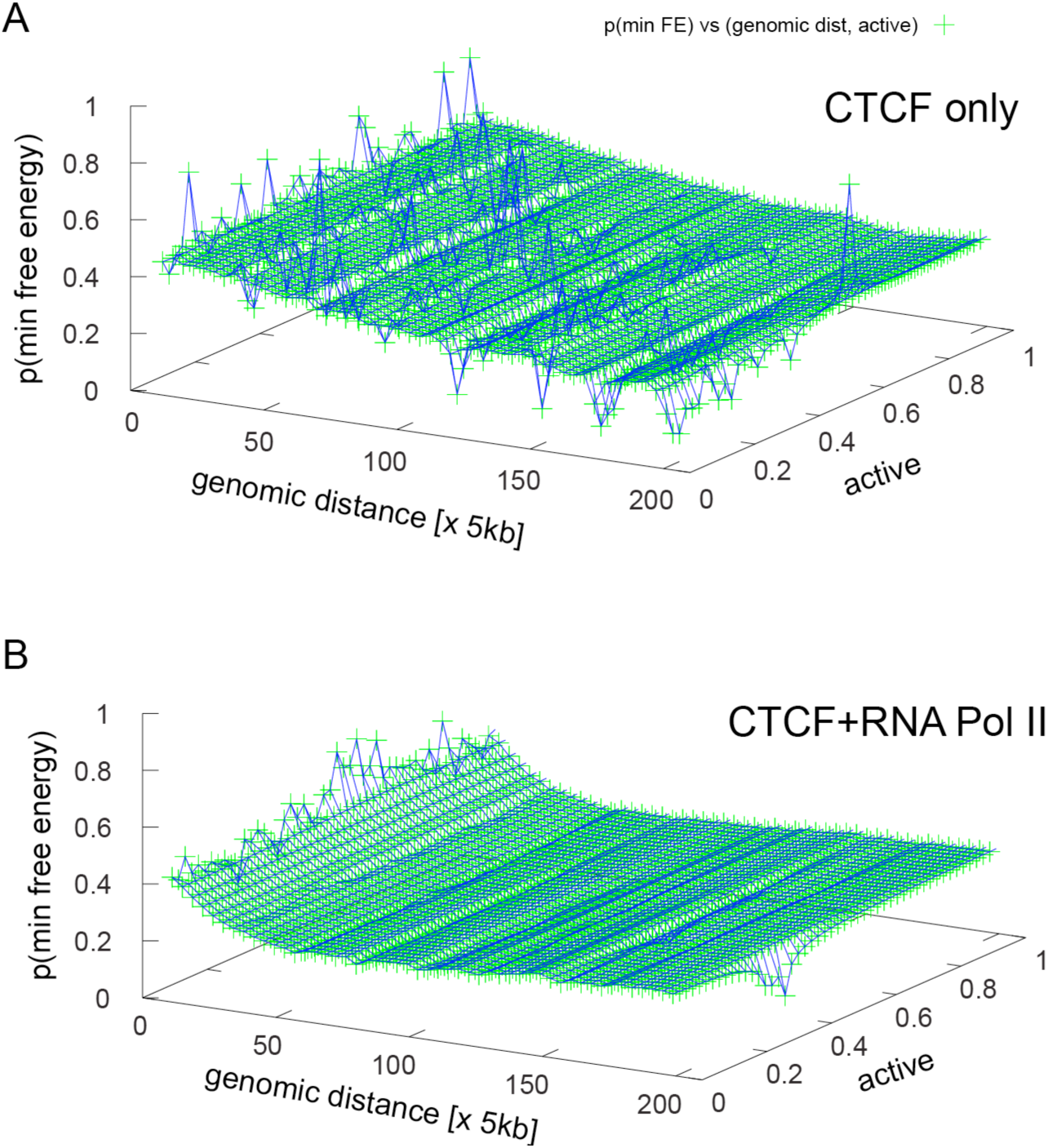
Comparison of the Boltzmann probability of the optimal structure plotted with respect to both the genomic distance and the degree of activity for CTCF (A) and RNA Pol II (B) loops. There does not appear to be a substantial difference in the probability as a function of activity, only with respect to genomic distance.

Figure 6 shows a plot of the free energy as a function of the genomic distance. There is a small trend in the data suggesting that the slope of the curve flattens out at some point. This would happen if there are limits on the size of the domain (at least for a single CTCF binding site). A rough estimate of the limit would be around 2 Mbp, based on the current data. In general, CCDs tend to have a limited range of about 3 Mbp. There are very long-range contacts that occur; however, these much larger regions are more likely to be due to contact with the nuclear membrane, a consequence of confinement inside the nucleus of the cell. It should be no surprise that the nuclear membrane can provide sufficient restraint, but is beyond the scope of this work. Likewise, the contacts of TADs can be adjoined through a tandem collection of pseudoknots: e.g., `“..A..B..a..C..b..D..c…E..d..e..”`, etc. Nevertheless, in all cases, each pair still remains under 3 Mbps.

**Figure 6.**
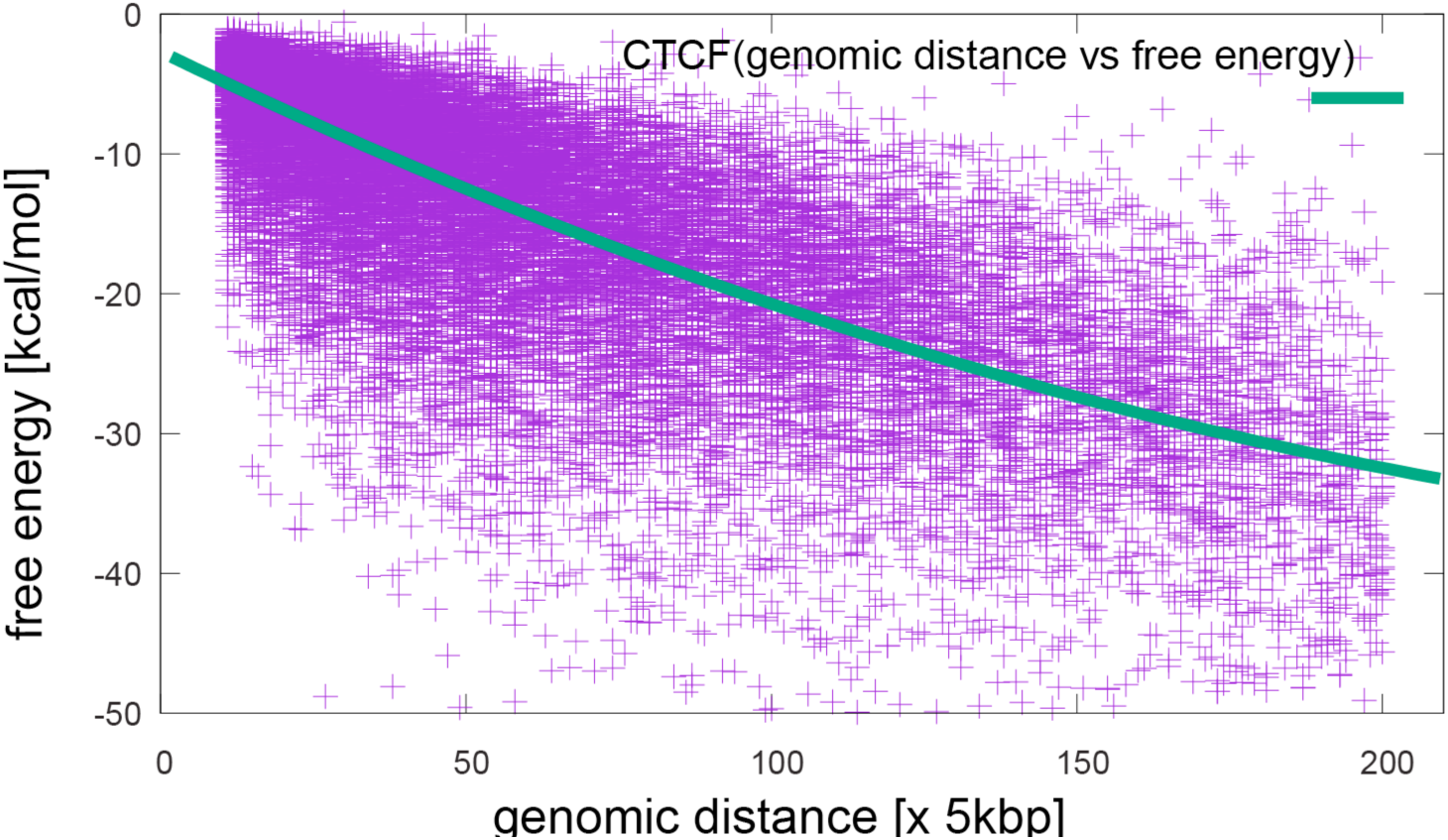
Plot of the free energy with respect to the genomic distance of the domain. The fit shows a slight quadratic component indicating that there may be a maximum in the size of the TAD.

Hence, we observe a model in which the mFE of CCDs as a function of genomic distance correlates and is consistent with active and open weight where inactive (closed) structures have the mFE. Moreover, the entropy estimates are largely correlated with the domain size for pairs in the TAD. Ensembles that have a dominant structure (a more negative free energy with large differences in the thermodynamic probability between the principal structure and suboptimal structures in the list) tend to be inactive on the one extreme. Ensembles with several very different conformations of nearly equal weight – a weaker free energy (FE) and small differences in the thermodynamic probability between very diverse structures - tend to be active structures.

Tied together, this suggests that the free energy evaluation strategy can aid in estimating epigenetic feature in the genome and the approximate structure of the observe contacts, or can incorporate these into computations. This work has also shown a proof of principle that an entropy based model used particularly in RNA structure prediction is also applicable in chromatin structure prediction.

For structures at 5 kbp resolution and 350 points, the calculations were able to run in a few minutes with roughly 5000 suboptimal structures produced. Several improvements would be advised. First, the simple pair weight function (Supplement Equation 3) is evidently far too simple because, whereas it appears to reproduce a lot of the heatmap, the intensities are not the same. This may also be reflecting issues with the ensemble of cells from which the pair weights were obtained. Perhaps single cell data would resolve this issue.

Similarly, it still appears that shaving off the dominant Boltzmann weights during the calculation may somewhat bias the ensemble set of structures that are reported. It may be better to calculate the full partition function and for building suboptimal structures, to select by domains rather than by total free energy.

## Conclusion

We have introduced a computational algorithm for estimating the structure and dynamics of chromatin loops by analyzing the thermodynamic probability of the minimum free energy structure. This permits learning the actual structure of the chromatin in terms of 2D structural motifs. Significant correlation was suggested by the tendency of known active structures to show a less stable minimum free energy (mFE), indicating that the observed structure of chromatin within the ensemble should be changing regularly with a high probability; likewise, the inactive regions tended to show structures with a very large mFE with respect to genomic distance. This algorithm may help serve as an aid in determining the relative activity of the chromatin based upon the stability of the structure. As with protein structure, the notion of a funnel shaped energy landscape originally proposed in protein folding (Bryngelson et al., 1995; Bryngelson and Wolynes, 1987; Onuchic and Wolynes, 2004) is also largely affirmed in chromatin structure as well (Zhang and Wolynes, 2017), at least at the meta-structural level studied here.

## Acknowledgements

DP, WD are supported by grants from the Polish National Science Centre (2014/15/B/ST6/05082 and 2013/09/B/NZ2/00121), the European Cooperation in Science and Technology action (COST BM1405 and BM1408). DP is supported by funds from National Leading Research Centre in Bialystok and the European Union under the European Social Fund. All authors were supported by grant 1U54DK107967-01 “Nucleome Positioning System for Spatiotemporal Genome Organization and Regulation” within 4DNucleome NIH program. Thanks to Przemyslav Szalaj for contributing the heat maps for different loops, Teresa Szczepinska for adding measures of activity and openness of the chromatin loops in the heat maps, and Michal Kadlof for providing advice on chromatin loops and providing the core python code for displaying 2D heat maps.

